# Comments on snails of the genus *Zoniferella* from Ecuador (Mollusca: Achatinidae), with restriction of the type locality “Los Puentes” for several species of Gastropoda and Arachnida

**DOI:** 10.1101/2022.03.03.482746

**Authors:** Diego F. Cisneros-Heredia, Roberto F. Valencia

## Abstract

The genus *Zoniferella* includes six taxa of land snails from Colombia and Ecuador for which little has been published beyond their original descriptions. In this paper, we present new records of *Zoniferella vespera*, a species previously known only from its type locality, expanding its range across northwestern Ecuador. We provide the first description of the colouration in life for *Zoniferella* snails. We comment on the similarities among some species of *Zoniferella*, suggesting the possibility that *Z. vespera* and *Z. riveti* are junior synonyms of *Z. albobalteata*, and *Z. bicingulata* is a synonym of *Z. riveti* var. *bizonalis* (as *Z. bizonalis*). Finally, we offer evidence to allow the restriction of the locality of “Los Puentes”, type locality of *Mesembrinis vesperus* Jousseaume, 1887 = *Zoniferella vespera* (Gastropoda: Achatinidae); *Isomeria bourcieri lutea* Cousin, 1887 (Gastropoda: Labyrinthidae); *Guestieria locardi* Jousseaume, 1887 (Gastropoda: Scolodontidae); *Proserpinella cousini* Jousseaume, 1887 (Gastropoda: Proserpinellidae); *Idiophthalma robusta* Simon, 1889 (Arachnida: Barychelidae); *Eurypelma* (*Lasiodora*) *augusti* Simon, 1889 = *Pamphobeteus augusti* (Arachnida: Theraphosidae), *Eurypelma* (*Lasiodora*) *vespertinus* Simon, 1889 = *Pamphobeteus vespertinus* (Arachnida: Theraphosidae).

---

*Zoniferella* Pilsbry, 1906 was described as a subgenus of *Synapterpes* Pilsbry, 1896, and later recognised as a distinct genus (Bank 2017, MolluscaBase 2022). *Zoniferella* are land snails characterised by a fragile, thin, glossy greenish-black shell, varying from yellowish olive green on the apex to dark greenish black on most of the shell, having white stripes. Size, number of whorls and stripes vary according to the species and soft anatomy remains unknown (Pilsbry 1906). Six taxa of *Zoniferella* have been described: *Zoniferella albobalteata* (Dunker, 1882) (type species by original designation), *Z. vespera* (Jousseaume, 1887), *Z. riveti* (Germain, 1907) and *Z. riveti* var. *bizonalis* (Germain, 1907), *Z. bicingulata* (Fulton, 1908), and *Z. pilsbryi* (Fulton, 1908). The latter was assigned to *Zoniferella* with uncertainty in its original description.

Nothing has been published in academic sources about *Zoniferella* species, aside from their original descriptions. This paper aims to present the first records for *Zoniferella vespera* since its original description, expanding its range across northwestern Ecuador, to comment on other species of *Zoniferella* from Ecuador, and to restrict the type locality of *Mesembrinus vesperus* Jousseaume, 1887 (= *Zoniferella vespera*), which is also the type locality of several other species of gastropods and spiders.

## *Zoniferella vespera* (Jousseaume, 1887)

*Mesembrinus vesperus* Jousseaume, 1887: 168, pl. 3, fig. 2

*Mesembrinus vesperus*—(Cousin 1887): 234; (Astudillo Espinosa 1978): 57.

*Bulimulus visendus*—(Hidalgo 1893b): 47–49; (Hidalgo 1893c): 247–248

*Bulimulus visendus* var. *vesperus*—(Hidalgo 1893a): 131

*Synapterpes* (*Zoniferella*) *vesperus*—(Pilsbry 1906): 234, pl. 37, fig. 91

*Synapterpes vesperus*—(Germain 1907): 61; (Germain 1911): C48.

*Zoniferella vespera*—(MolluscaBase 2022): AphiaID 995660

Two individuals of *Zoniferella vespera* were found at the Otonga Reserve (0°26’24” S, 78°43’19” W; 1815 m), county of Sigchos, province of Cotopaxi, Republic of Ecuador, on top of leaves of Melastomataceae and Araceae, in a very narrow, steep, and rocky ravine covered by old-growth dense forest, near the main station house, on 28 May 2021 by Roberto Valencia (Fig. 1). The snails were photographed alive, and one is deposited at the Museo de Zoología, Universidad San Francisco de Quito, Ecuador (ZSFQ), under patent 010-UBVS-OTQ-DZ2E-MAAE-2021, issued by the Ministry of Environment, Water and Ecological Transition of Ecuador (Fig. 2).

**Figure 1.**
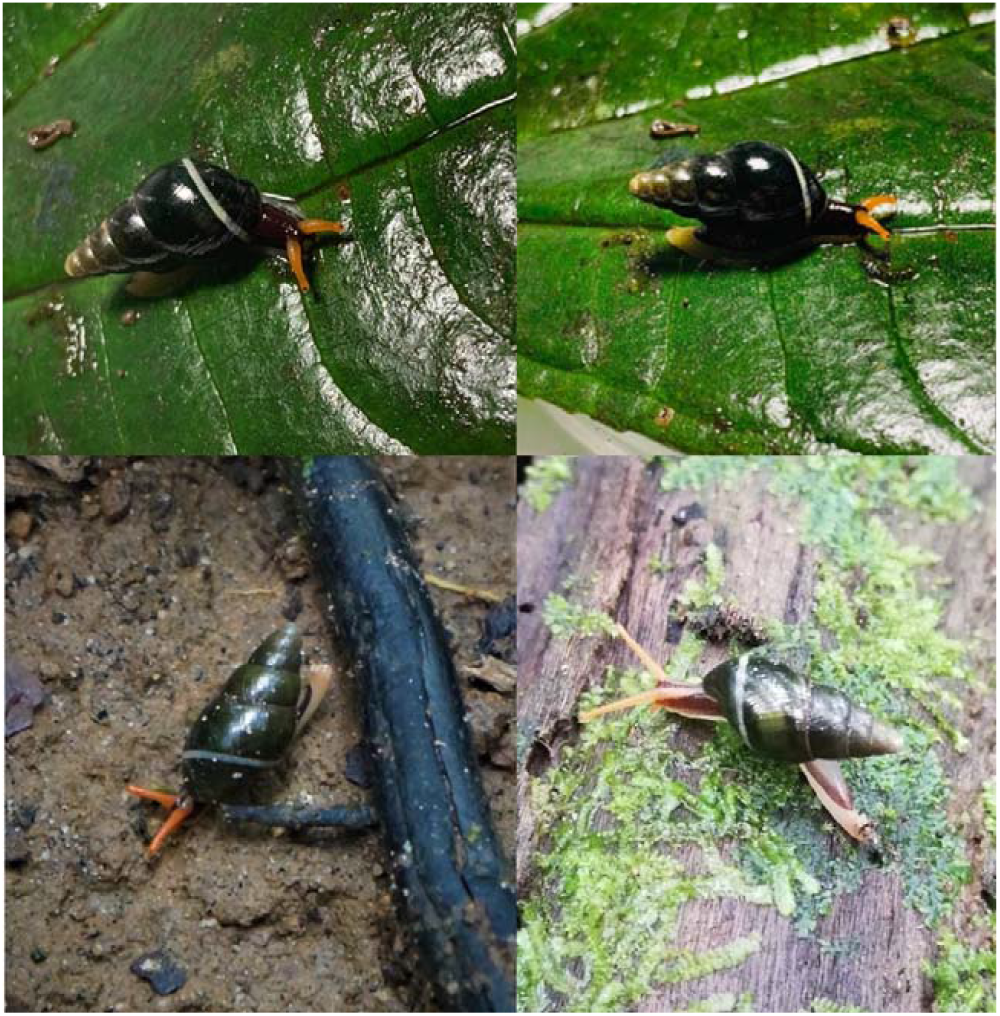
Variation of colouration in life of *Zoniferella vespera*. Top: Otonga Reserve, province of Cotopaxi, Ecuador, photo by R. Valencia. Bottom left: Un Poco de Chocó Reserva, Pachijal, province of Pichincha, Ecuador, photo by j0rdis, iNaturalist (CC-BY-NC). Bottom right: Estación Mindo USFQ, province of Pichincha, Ecuador, photo by Giovanni Ramón, iNaturalist (CC-BY-NC-SA)

**Figure 2.**
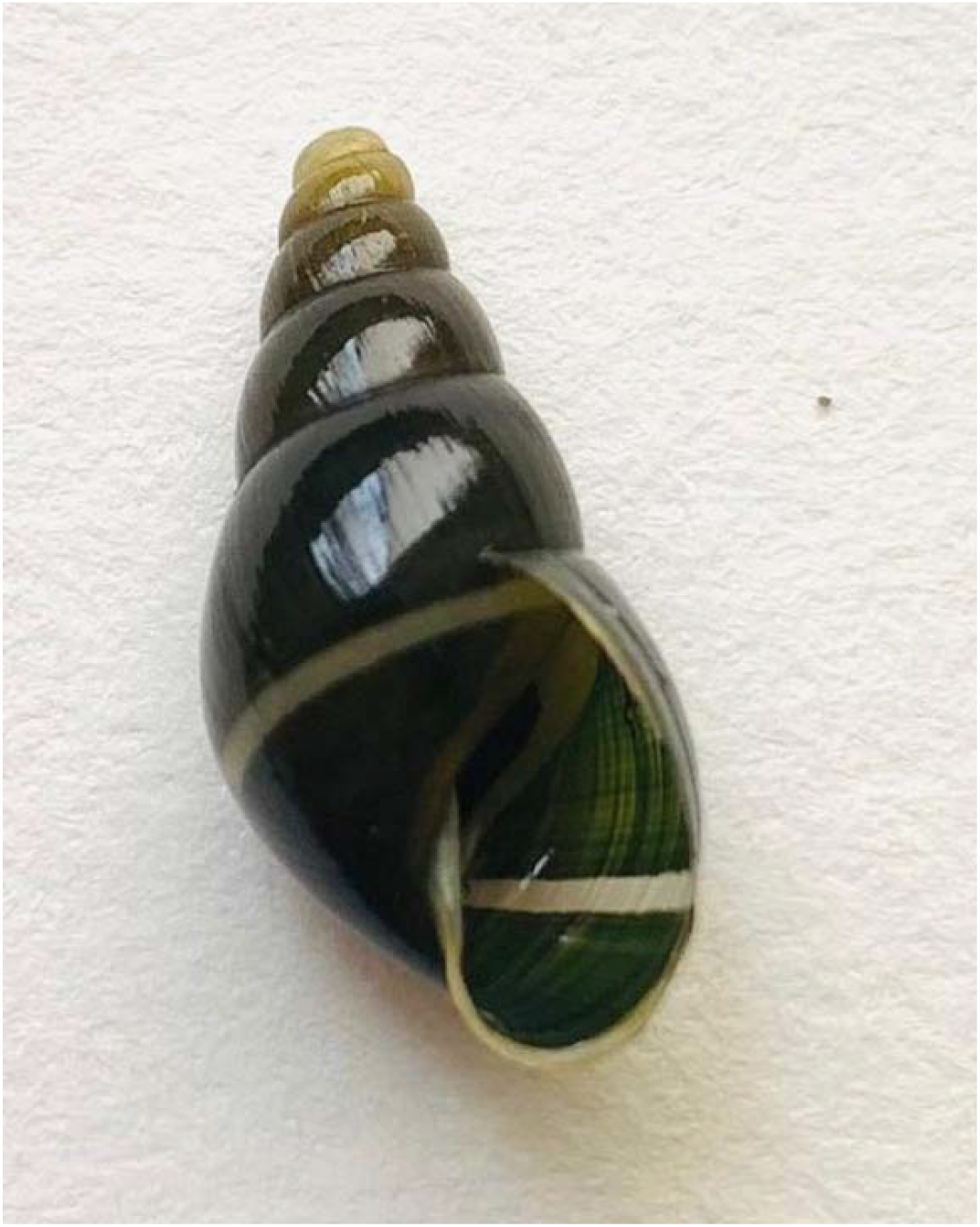
Specimen of *Zoniferella vespera* from Otonga Reserve, province of Cotopaxi, Ecuador, deposited at the Museo de Zoología, Universidad San Francisco de Quito (ZSFQ).

Photographs of one individual were posted in iNaturalist (Valencia 2021), where initial generic identification was provided. We confirmed the generic and species identification based on information provided by Pilsbry (1906), Jousseaume (1887) and Cousin (1887), and examination of photographs of the holotype (MNHN 2000b). Snails were identified as *Zoniferella vespera* due to the presence of the following diagnostic characters: shell ovate-acuminate, thin, delicate, very glossy, decorated with very thin irregular ridges; 6 whorls; spire long, conic; apex obtuse; aperture oval; peristome simple, acute; columellar margin straight. Shell colouration dark green getting paler towards the apex, with a circular white conspicuous band on the last whorl, and columellar margin whitish (Cousin 1887, Jousseaume 1887, Pilsbry 1906). The new specimen is slightly more globular and smaller than the holotype, but we consider that these differences are due to intraspecific variation (Fig. 2).

*Zoniferella vespera* is most similar to *Z. albobalteata* and *Z. riveti. Zoniferella albobalteata* is currently diagnosed from *Z. vespera* by its smaller length (13 mm vs. 17 mm in the holotype of *Z. vespera*) and subreflexed columella (straight in *Z. vespera*) (Dunker 1882, Pilsbry 1906). However, size of snails from Otonga is intermediate between both species, but due to the straight columella we assigned them to *Z. vespera. Zoniferella riveti* differ from *Z. vespera* by having 7 whorls, a larger size (21 mm), more slender shape, and twisted columella (Germain 1907, 1911). *Zoniferella riveti* var. *bizonalis* and *Z. bicingulata* differ from *Z. vespera* by having two white bands on the last whorl, one of which further continues along the other whorls (one band in *Z. vespera*, restricted to the last whorl) (Germain 1907, 1911, Fulton 1908). *Zoniferella pilsbry* is easily separated from *Z. vespera* by having the lower whorls with narrow spiral bands green alternating with narrow white bands, eight whorls, and longer length (26 mm) (Pilsbry 1906).

Subtle differences in length (13–21 mm), number of whorls (6–7) and columella shape (straight or twisted) have been used to separate *Z. albobateata, Z. vespera* and *Z. riveti*. A similar situation is noted between *Z. riveti* var. *bizonalis* and *Z. bicingulata*, and both species were not compared in their original descriptions because they were published a year apart (Germain 1907, Fulton 1908). Based on the data provided in their descriptions and photographs of the holotypes (MNHN 2000a, NHM 2021), they would differ in the number of whorls (6½–7½) and the shape of the second white band (narrower than the first band in *Z. r. bizonalis*, and the same size in *Z. bicingulata*). However, these differences could be intraspecific variation of single species, suggesting the possibility that *Z. vespera* and *Z. riveti* are junior synonyms of *Z. albobalteata*, as suggested by Hidalgo (1893c), and *Z. bicingulata* a synonym of *Z. riveti* var. *bizonalis* (as *Z. bizonalis*). *Zoniferella albobalteata* is currently known only from its type locality in “humid forests near Pasto, Colombia” (Dunker 1882, Pilsbry 1906), about 180 km N from the known range of *Z. vespera*, while the type locality of *Z. riveti* (San Tadeo hill, road to Pachijal) is about 10 km W from the type locality of *Z. vespera*, and within its distribution range (see below). Unfortunately, the type locality of *Z. bicingulata* was not specified, cited only as Ecuador (Fulton 1908). Decisions about their formal synonymy would require the direct comparison of type specimens and fresh topotypic material.

The colouration in life of *Z. vespera* has not been described. Individual of *Z. vespera* from the Otonga Reserve had the sole and border of foot yellowish cream, dorsal surfaces of foot and head dark purplish, and tentacles bright orange (Fig. 1). This colouration in life is observed in other individuals reported in iNaturalist, with some intraspecific variation observed by some individuals having a whitish or orange band running towards the back from the base to each tentacle, and the tentacles coloured yellowish cream as the border of foot (Fig. 1). The colouration in life of the shell is like the coloration in preservation, but some individuals may look almost black. Photographs reported in iNaturalist of individuals with two bands on the last whorl (*Z. riveti* var. *bizonalis* / *Z. bicingulata*) usually differ by having the tentacles completely dark like the dorsal surfaces of foot and head, except for the eyes that are cream (Fig. 3). However, one individual showing two bands has the same colouration described for *Z. vespera* (Obando 2020) and another has tentacles completely cream-coloured and a whitish band running towards the back from the base to each tentacle (Stegenga 2022) (Fig. 3).

**Figure 3.**
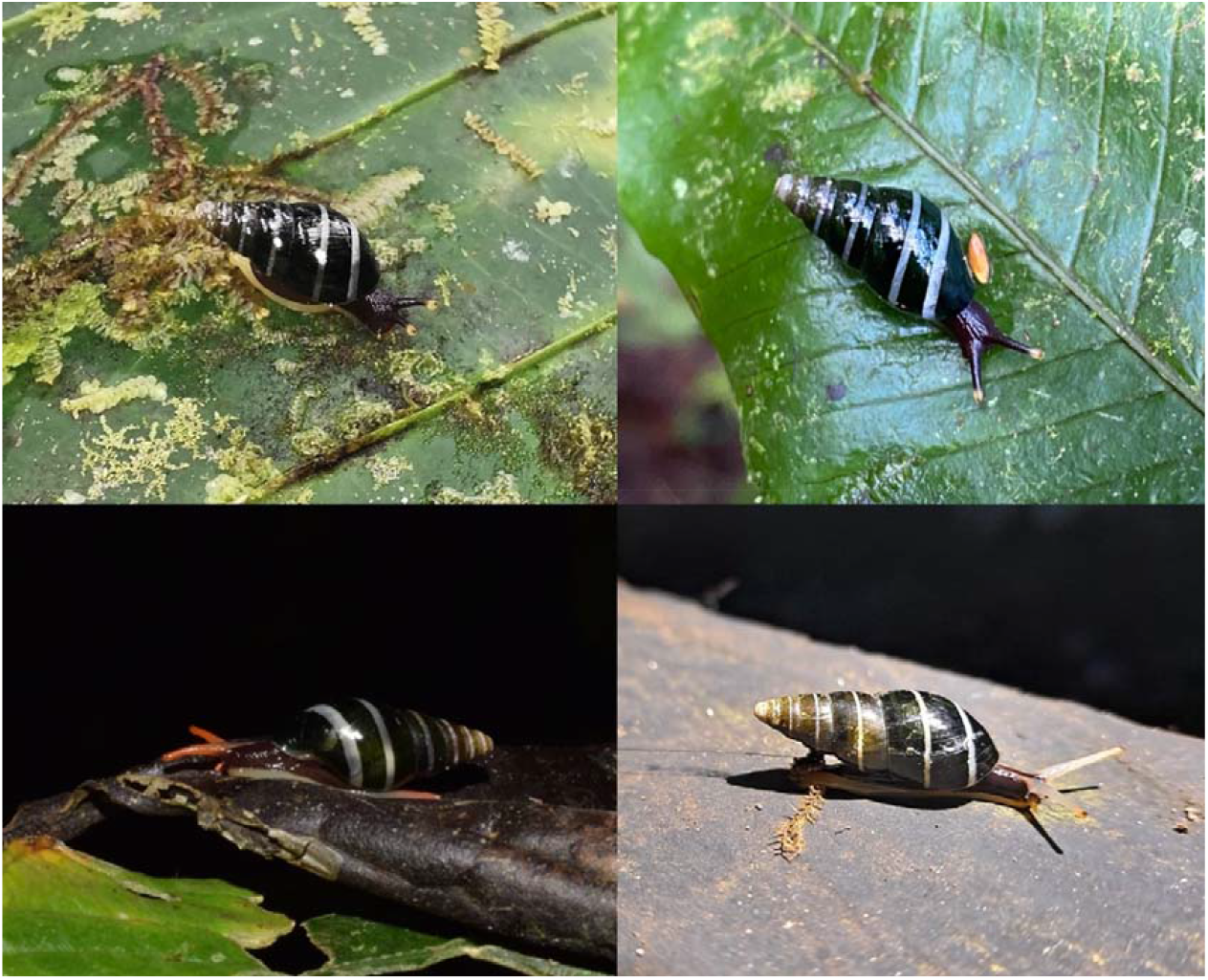
Variation of colouration in life of *Zoniferella riveti* var. *bizonalis* / *Z. bicingulata*. Photos by Barna Takats (top left), Seth Ames (top right), Ben Stegenga (bottom left), Eduardo Obando (bottom right), iNaturalist (CC-BY-NC).

“Los Puentes” is the type locality of several taxa of molluscs and spiders, including *Zoniferella vespera* (Jousseaume, 1887); *Isomeria bourcieri lutea* Cousin, 1887; *Guestieria locardi* Jousseaume, 1887; *Proserpinella cousin* Jousseaume, 1887; *Idiophthalma robusta* Simon, 1889; *Pamphobeteus augusti* (Simon, 1889); and *Pamphobeteus vespertinus* (Simon 1889) (Cousin 1887, Jousseaume 1887, Simon 1889, Breure 2020). However, the precise position of “Los Puentes” has not been established and some authors became confused by descriptions stating that it was “near Quito”, suggesting that Los Puentes was in the inter-Andean valley of Quito. Jousseaume (1887) provided the following information about the type locality of *Mesembrinus vesperus* (now *Zoniferella vespera*): “A single example of this species was collected by our colleague, Mr. A. Cousin, in his property of Los Puentes, near Quito”. Simon (1889) cited the type locality of *Idiophthalma robusta* as *“*Los Puentes, near Quito”; he reported specimens of *Linothele longicauda* (Ausserer, 1871) and *Cyclosternum schmardae* Ausserer, 1871—whose type locality is Quito, from *“*Los Puentes, near Quito” and “Los Puentes…, around Quito”. For *C. schmardae*, Simon (1889) said “we also received it from Los Puentes and Rumipamba, around Quito, by Mr. A. Cousin”, implying that, like Rumipamba, Los Puentes was in the inter-Andean valley of Quito.

Most collections from Los Puentes were obtained by Auguste Edouard Cousin (Paris, 1835– Quito, 1899, see Breure 2020, Correoso Rodríguez 2020). Cousin published a single but important academic publication: “*Faune malacologique de la République de l’Équateur*” (Cousin 1887), where he regularly cited “Los Puentes, near Gualea” when commenting on the distribution of several species (e.g., *Cyclophorus esmeraldensis, Mesembrinus visendus, Porphyrobaphe irroratus, Drymaeus petasites, Ammonoceras flora, Psadara selenostoma, Isomeria bourcieri*, and *Obeliscus cuneus*), and said that he found *Bourciera helicinaeformis* “On the road from Quito to Gualea, towards Hacienda de Los Puentes”. Cousin cited “Los Puentes, parish of Calacalí, prov. of Pichincha” in the distribution of *Eurytus taylorianus*, and “Los Puentes, county of Quito” for *Isomeria cymatodes* and *Cyclotus fischeri*. Cousin also provided information about the altitude of Los Puentes, saying that it was “More of less 1500 m” for *M. visendus* and “about 1500 m” for *O. cuneus*. Pilsbry (1906) cited the locality as “Los Puentes near Gualea at about 1500 meters… (Cousin)” for *O. cuneus*, based on the information provided by Cousin (1887). Paul Rivet apparently collected at Los Puentes during his expeditions of the Second Geodesic Mission to Ecuador, since Germain (1907, 1911) cited specimens of *Bulimus* (*Porphyrobaphe*) *irroratus* collected by Rivet at “Los Puentes, road of Gualea”. Francisco Cousin, son of Auguste Cousin, was friend with Rivet (Jarrín 2021), and probably invited Rivet to the Cousin family’s farm at Los Puentes. Breure (2020) reported that the label accompanying the types of *Proserpinella cousini* said “Los Puentes San Fernandino”, however, the photographs of the labels shown that it was written “Los Puentes S. Fernando” and not “Fernandino” (Breure 2020: figs. 31–32). Correoso Rodríguez (2020) mentioned that “Los Puentes” was in northwestern Pichincha and that Auguste Cousin bought the farm in 1866.

Based on the map by Wolf (1892), a bridle path connecting Quito with Gualea started going north across the inter-Andean valley of Quito towards the towns of Cotocollao and Pomasqui, then climbing the western Andean Mountain range through the Casitagua and Pululahua mountain pass towards Calacalí and Nono and running down the western slopes of the Andes towards Nanegal and Gualea (Fig. 4). Los Puentes is mentioned in an executive decree issued by Eloy Alfaro, president of Ecuador, on 14 June 1898 and included in the report presented by Ricardo Valdivieso of the Ministry of Public Works to the Congress of 1899 (Valdivieso 1898). The decree responded to requests submitted by people from the region of Gualea to improve the local bridle path and confirms that the farm Los Puentes was in the parish of Nanegal and on the Quito–Nono–Gualea road:

**Figure 4.**
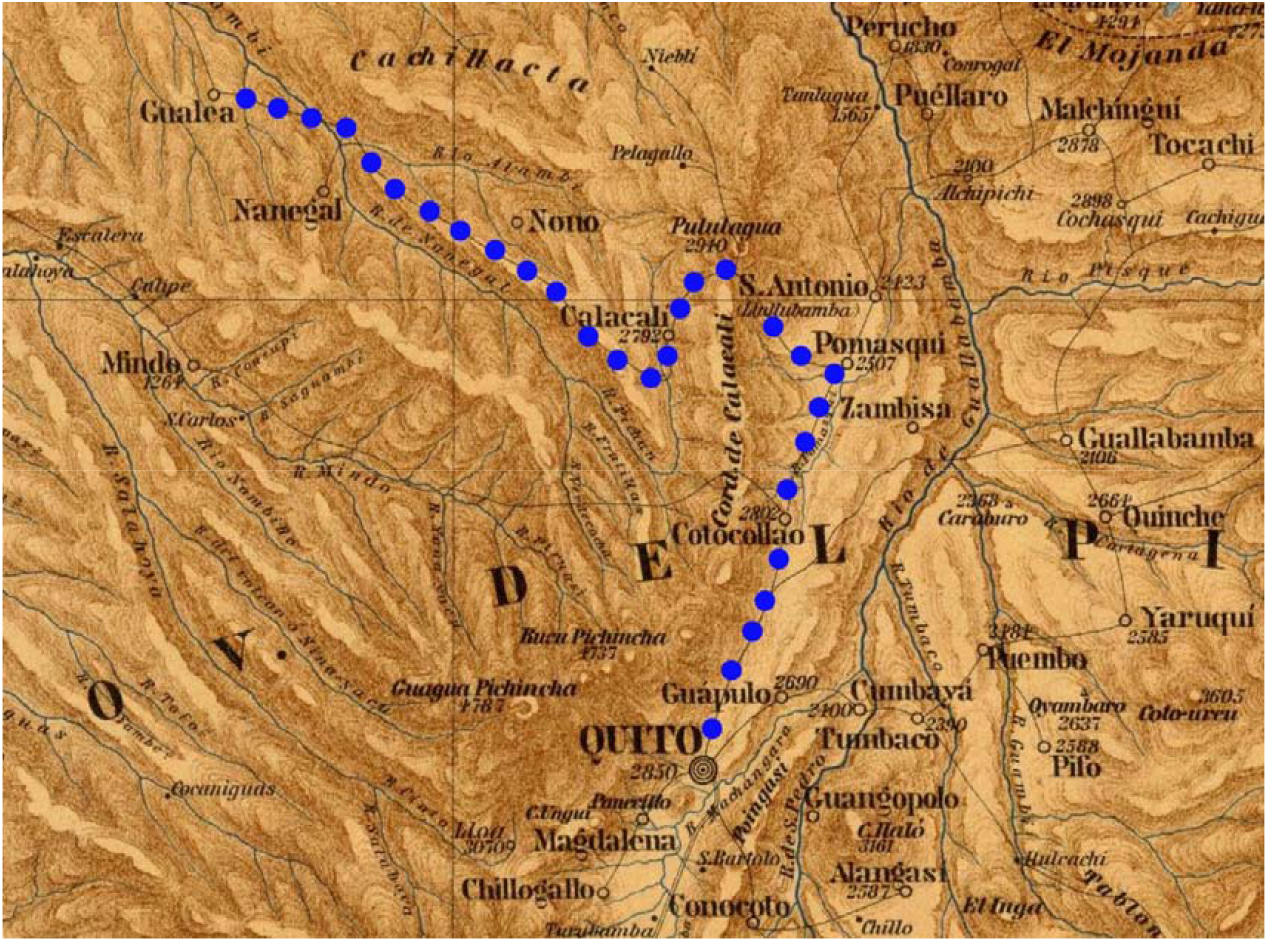
Section of the Carta Geográfica del Ecuador by Theodore Wolf (1892) indicating the Quito–Gualea bridle path (blue dots). Dark lines indicate roads and bridle paths.

Translation: Art. 1° Repair the bridle path from Gualea to Nono. Art. 2° For this work the owners of rustic estates of the parish of Gualea will contribute, once only, with 7% of the value of their properties. The farms “Chiquilpe” of the parish of Nono and “Los Puentes” of Nanegal, will contribute with the same contribution.

During the second half of the 20^th^ century, sections of the Quito–Nono–Gualea bridle path were the basis for the construction of the Quito–Nono–Nanegalito road and the Calacali-Nanegalito-Puerto Quito highway. “Los Puentes” literally means “the bridges” in Spanish, and there were many bridges in the old bridle path from Quito to Gualea in the 19^th^ century. However, there is a single locality called “Los Puentes” in the region of the old bridle path and modern road/highway, nowadays classified as a neighbourhood of the parish of Nanegal (IGM 1990, DIPLA n.d., GAD Pichincha n.d.). This locality lies very close to an area where large bridges have been historically built to cross a river junction. About 2 km NE from this locality, there is an area known as “San Fernando Cuatro Hermanos”, which would coincide with the locality mentioned for *Proserpinella cousini*.

The correct definition of a type locality is key to understand the distribution of a species because it corresponds to the original locality where the name-bearing type was collected. Following recommendation by (ICZN 1999, see discussion by Cisneros-Heredia 2017), we restrict the type localities of *Isomeria bourcieri lutea* Cousin, 1887 (Gastropoda: Labyrinthidae); *Guestieria locardi* Jousseaume, 1887 (Gastropoda: Scolodontidae); *Proserpinella cousini* Jousseaume, 1887 (Gastropoda: Proserpinellidae); *Mesembrinis vesperus* Jousseaume, 1887 (Gastropoda: Achatinidae); *Idiophthalma robusta* Simon, 1889 (Arachnida: Barychelidae); *Eurypelma* (*Lasiodora*) *augusti* Simon, 1889 (Arachnida: Theraphosidae), *Eurypelma* (*Lasiodora*) *vespertinus* Simon, 1889 (Arachnida: Theraphosidae), and any other species whose name-bearing type material was collected at “Los Puentes”, the farm of the Cousin family in northwestern Ecuador, as follows: Los Puentes, near Nanegalito, at 1500 m elevation, province of Pichincha, República del Ecuador.

Coordinates focus point: 0.045833, -78.674167, radius: 2 km. The newly restricted type locality is described as a circle, with a focus point and a radius to describe the associated uncertainty as a maximum distance from that point within which the locality is expected to be found (i.e., point-radius method, Wieczorek et al. 2004). A KMZ file showing this locality is available here: https://doi.org/10.6084/m9.figshare.19297532. In addition to Los Puentes and Otonga Reserve, observations of *Z. vespera* have been reported on iNaturalist from several localities in northwestern Ecuador: valley of Mindo and surrounding mountains, Pachijal, Mashpi, province of Pichincha; and, San Francisco de las Pampas, province of Cotopaxi (Fig. 5, see Supplementary file 1).

**Figure 5.**
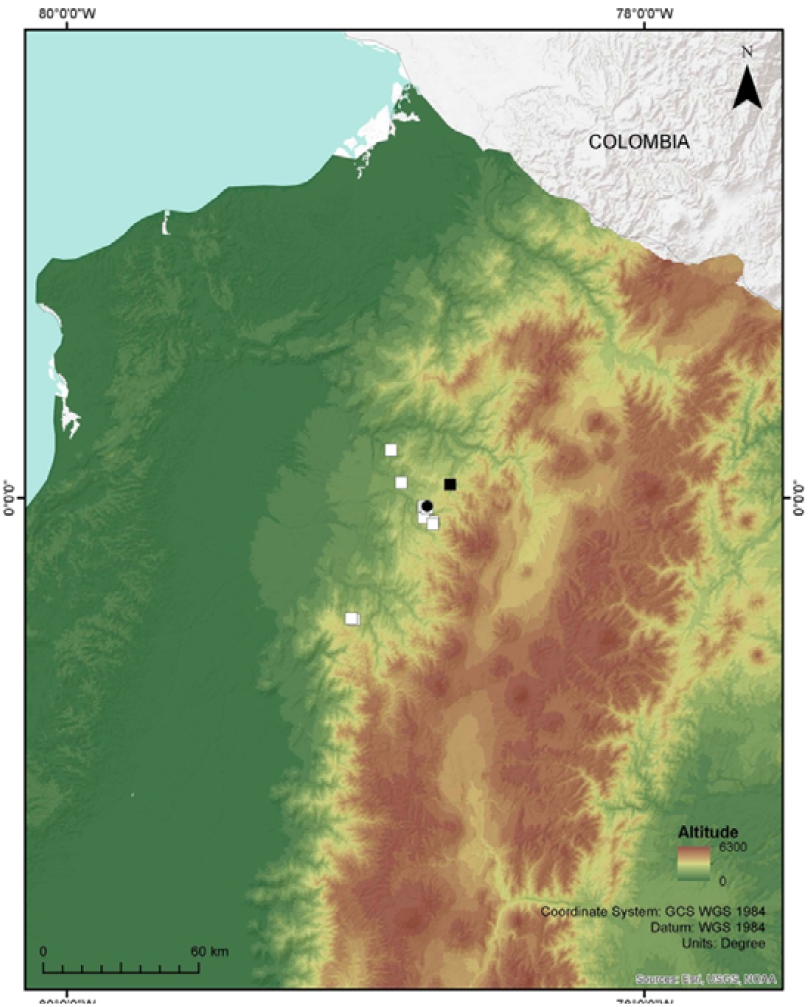
Map of northwestern Ecuador showing the type locality (black square) and other known localities (white squares) of *Zoniferella vespera*, and the type locality of *Z. riveti* and *Z. riveti* var. *bizonalis* (black circle).

Extensive habitat changes and loss across Ecuador have most probably threatened and pushed towards extinction several snail species. The montane cloud forests on the northwestern Andean slopes of Ecuador, habitat of *Zoniferella vespera*, have been deeply affected by deforestation for timber extraction, agricultural expansion, and mining projects, with few old-growth forest fragments remaining in the region. Information on the diversity, ecology, and biogeography of terrestrial snails of Ecuador is deficient. Aside from some works by Abraham Breure, Francisco Borrero, Modesto Correoso and collaborators (Breure and Borrero 2008, Correoso Rodríguez 2008, 2010, Borrero and Araujo 2012, Breure and Araujo 2017, Breure 2020), little has been published on the fauna of terrestrial snails from Ecuador in recent years.

## Acknowledgements

We thank Sergio Norona, guide at Otonga Reserve, and Roberto León and Samuel Cortese for their help in the field; Giovani Ramón and Emilia Peñaherrera of the Museo de Zoología of Universidad San Francisco de Quito USFQ for their support during lab work; Matt Parr for suggesting the generic identification in iNaturalist; and xx reviewers for their comments on the manuscript. We are grateful to the Otonga Reserve for allowing us to venture into the cloud forest that they have protected for decades; to the Natural History Museum London and the Muséum national d’Histoire naturelle for offering online photographs of type specimens; and to the Biodiversity Heritage Library BHL, Google Books, Archive.org and the World Spider Catalog for making important literature freely available. This work was supported by Universidad San Francisco de Quito USFQ through research and outreach funds (HUBI ID 1057, 607) and operative funds assigned to IBIOTROP.

---

**Supplementary file 1.**
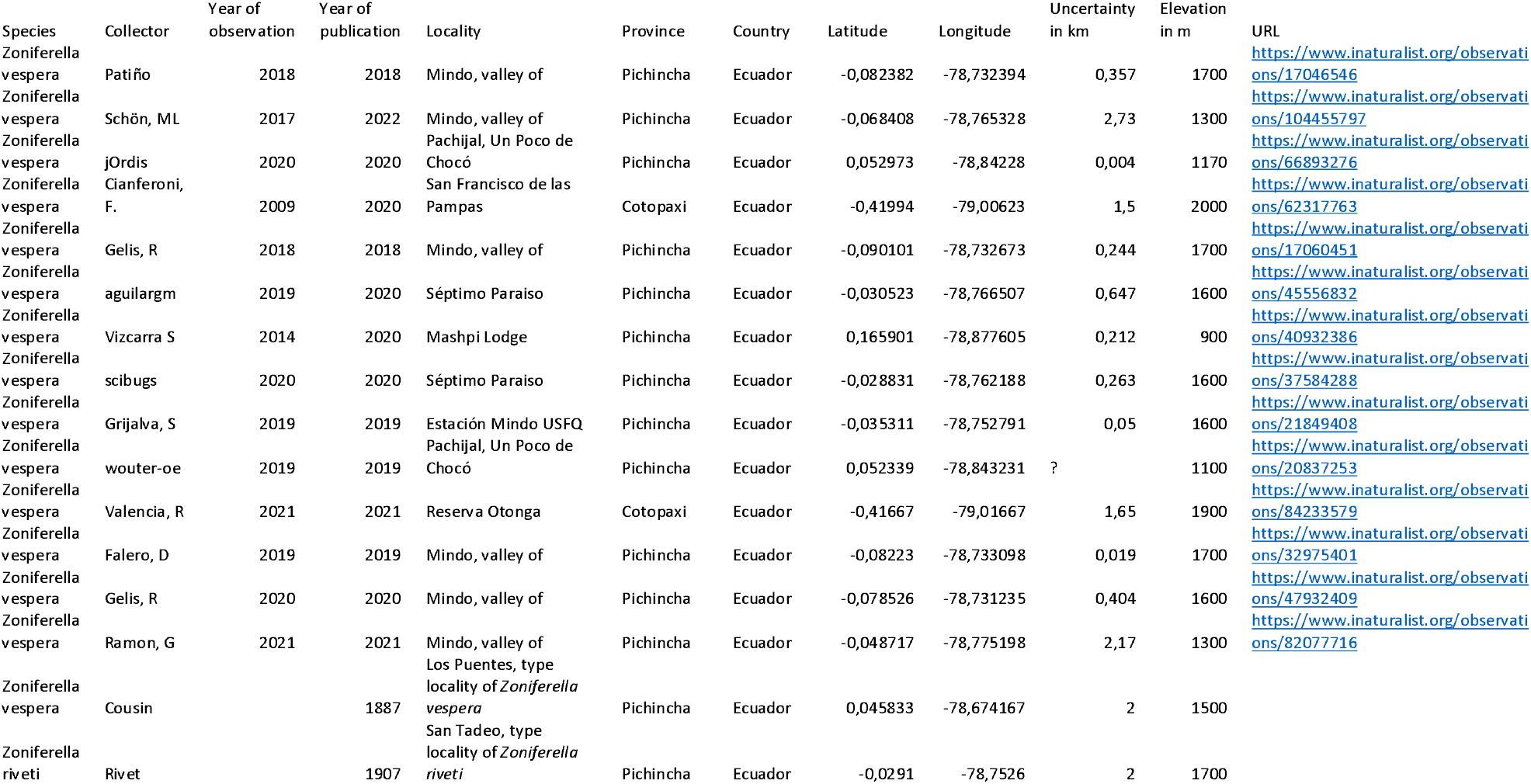
Known localities of *Zoniferella vespera* (Jousseaume, 1887) and *Z. riveti*.

